# Development of early-stage type 1 diabetes in germ-free interleukin-10 deficient mice

**DOI:** 10.1101/2020.04.18.048272

**Authors:** Alexandria M. Bobe, Jun Miyoshi, Patrick Moore, Suzanne Devkota, Vanessa Leone, Kristina Martinez, Betty R. Theriault, Mark W. Musch, Clive Wasserfall, Mark Atkinson, Christopher J. Rhodes, Eugene B. Chang

## Abstract

Several experimental models demonstrate a role for gut microbiota in the progression of type 1 diabetes (T1D) in genetically prone hosts. While the association between disturbances in gut microbiota, or microbial dysbiosis, and complex immune diseases such as inflammatory bowel diseases (IBD) are well established, less is known about its role in T1D pathogenesis. In IBD-prone interleukin-10 deficient (IL-10 KO) mice, the absence of gut microbiota under germ-free (GF) conditions prevents IBD development. However, in aged GF IL-10 KO mice (>6-months of age), polyuria and pancreatic lymphocytic infiltration resembling T1D lesions was observed. Approximately 50% of male and female mice above 6-months of age develop pancreatic immune cell infiltration, as compared to none in conventionally-raised and fecal microbiota transplanted (FMT) IL-10 KO counterparts. Immunofluorescence staining of islet infiltrates was positive for adaptive and innate immunological markers, including lymphoid and myeloid cell markers, which typically characterize autoimmune T1D lesions. A subset of GF IL-10 KO mice was also positive for insulin autoantibodies (IAA), but the majority of mice did not become diabetic. Our findings of early stage lymphocytic infiltrates in the pancreas and IAA in the absence of overt diabetes in GF IL-10 KO mice embody the early stages of T1D pathogenesis. As such, we propose that the presence of gut microbiota play a protective role against immune infiltration in the pancreas of genetically prone hosts. Moreover, our model provides an opportunity to better understand the role of the microbiota in the early stages of immune pathogenesis and perhaps conceive the development of microbe-mediated prophylactic strategies to treat or even prevent T1D.

---

The rise in T1D incidence over the last half century reflects the significance of environmental, societal, and dietary factors in disease pathogenesis. The heritability of T1D is estimated to be 72-88%, with triggering environmental factors thought to significantly contribute to the phenotypic variability noted in patients (Hyttinen *et al*., 2003). The pathophysiology of T1D is multifactorial and characterized by specific autoimmune destruction of pancreatic β-cells. Consequently, T1D patients become insulin therapy dependent. The immune cell infiltration of the pancreas reveals a prominent role for auto-reactive T cells in disease pathogenesis, along with antigen presenting cells and ineffective regulatory T cells (Tregs) (Tritt *et al*., 2008; Phillips *et al*., 2009; Tai, Wong and Wen, 2016). While progress has been made in identifying key players involved in islet invasion and β-cell destruction (S. Kim et al., 2007; Coppieters *et al*., 2012; Magnuson *et al*., 2014; Lundberg *et al*., 2017), prophylaxis and the precise etiology in humans still remains relatively elusive.

Several studies propose environmental triggers including viral infections, dietary components, antibiotic use and more recently, gut microbial dysbiosis, as contributing factors to disease onset and progression (Mejía-León and Barca, 2015; Hu, Wong and Wen, 2017). Over time, these environmental triggers may fuel metabolic and immune dysfunction that trigger “β-cell suicide or homicide”(Bottazzo, 1986; Atkinson *et al*., 2011). Distinct gut microbial profiles have been associated with T1D patients as compared to healthy controls (Dunne *et al*., 2014; Asnicar *et al*., 2017; Pellegrini *et al*., 2017). For example, T1D patients have been shown to display an “autoimmune microbiota” defined by decreased intestinal microbial diversity and a low abundance of short chain fatty acid (SCFA)-producing bacteria (Giongo *et al*., 2011). However, exact causation is difficult to discern from human studies and the gut microbial dysbiosis noted in these patients may be secondary to an already altered metabolic state in patients. Thus, mouse models provide a valuable opportunity to test variables in a controlled setting and dissect underlying mechanisms of disease pathogenesis that may translate to patients.

While our understanding of these complex interactions advances, additional models, particularly at the earliest presymptomatic stages of disease development, are needed to identify and optimize effective prevention and treatment strategies for patients at various stages in disease progression (Atkinson *et al*., 2011; Insel *et al*., 2015). The discovery of pancreatic inflammation in GF IL-10 KO mice creates a new model for understanding the role of the gut microbiome and anti-inflammatory cytokine, interleukin-10 (IL-10) in the early phases of immune initiation and disease progression. We employed the IBD-prone IL-10 KO mouse as a tool to understand the role of the gut microbiome in the development of T1D under GF conditions. Our initial characterization demonstrates GF IL-10 KO mice exhibit early stage T1D, with a small subset of mice progressing to overt diabetes (Bobe *et al*., 2017).

## Research Design and Methods

### Animals

Murine experimental procedures were performed at the University of Chicago and approved by The Institutional Animal Care and Use Committee (IACUC). Male and female C57Bl/6 Interleukin-10 deficient (IL-10 KO), wild-type (WT), and recombination-activating gene 1 deficient (RAG1 KO) mice were bred in-house, fed ad libitum, and maintained under a normal 12-hour light/dark cycle. Germ-free (GF) cohorts were group-housed and maintained on autoclaved chow (LabDiet, 5K67) in flexible film isolators in the University of Chicago Gnotobiotic Research Animal Facility (GRAF). Specific pathogen free (SPF) cohorts were group-housed and maintained on irradiated chow (Envigo, Teklad 2918 diet) in the same SPF vivarium room.

### Histological assessment

Harvested pancreata were OCT-embedded and frozen in a methyl-butane bath, or paraformaldehyde (4%) fixed at 4°C for 4-6 hours, processed, and paraffin embedded. Serial sections, 5-μm thick, were stained with immunofluorescence or Hematoxylin/Eosin (H&E). Samples were processed and sectioned at the University’s Human Tissue Resource Center (HTRC) or in-lab following HTRC’s protocol guidelines.

### Pancreatic infiltration incidence

To determine pancreatic infiltration incidence, whole H&E stained pancreas sections were scanned and scored for the presence or absence of lymphocytic infiltrates on a blinded basis in Panoramic Viewer. The presence of insulitis included peri-insulitis to mild or severe insulitis. This scoring was used to determine the incidence of pancreatic infiltration by calculating the number of mice positive for pancreatic immune cell infiltrates divided by the total number of mice scored in each mouse group.

### Immunohistochemistry

Immunostaining was performed as previously described (Alarcon, Verchere and Rhodes, 2012). To identify immune cell types within pancreatic infiltrates, frozen sections of pancreas were stained with primary antibodies: insulin (Sigma-Aldrich, MO); CD3 (Sigma); MHC-1 (Abcam, CA); CD4 (Abcam); CD68 (Abcam); B220 (BD Pharm, CA); CD8 (Thermo Fisher, MA); CD11c (StemCell Tech, Canada); and pSTAT1 (BD Bioscience, CA). Donkey derived secondary antibodies (Jackson ImmunoResearch) were used for fluorescence visualization on a DSU confocal microscope.

### Diabetes assessment

To assess diabetes development, several procedures were performed to track blood glucose levels in gnotobiotic isolators and the animal vivarium. To initially track blood glucose measurements of mice housed in isolators, submandibular cheek bleeds were performed by trained technicians in the GRAF unit and serum was stored at −80°C. Mice were monitored for increased frequency of urination and urinalysis was performed to test for glucose in the urine (Bayer, Keto-diastix Reagent Strips). Blood glucose levels (mg/dL) were recorded with a glucometer (Accu-Chek Compact Plus) from blood droplets from individual tails of mice (1) sacrificed immediately after 4-6 hours of fasting started in the morning (“Fasted Blood Glucose” reading), or (2) non-fasting mice sacrificed in the morning (“A.M. Blood Glucose” reading). Glucose tolerance tests were performed on mice fasted overnight for no more than 15-hours. Mice received intraperitoneal injections (IP) of 20% dextrose solution. Blood glucose readings were measured at 0, 15, 30, 60, and 120 minutes.

### Adoptive transfer

GF RAG1 KO mice recipients (15 males, 12 females ranging from 8-14 weeks of age) were IP injected with splenic immune cells (splenocytes) isolated from aged GF IL-10 KO donors (4 males, 4 females ranging from 27-35 weeks of age) under sterile conditions. One older GF WT male and female were used as control GF WT donors. Each adoptive transfer was performed with a donor to recipient ratio of 1:2 or 1:3. Fecal aerobic and anaerobic cultures were collected throughout the study and 16s rRNA gene PCR was performed to ensure the absence of microbial contamination in the isolator. GF RAG1 KO recipients were weighed weekly and removed from the isolator after 10-weeks for metabolic and histological assessment.

### Flow cytometry

Spleens were mashed through a sterile 70μm filter, washed with sterile filtered CRPMI (RPMI + 10% FCS) and spun down at 4C 600G for 5 minutes. Red blood cells were lysed (RBL Buffer: H_2_O 100ml, NH_4_Cl 0.824g, KHCO_3_ 0.1g, 0.5M EDTA 20uL), cells were spun down again and resuspended in sterile FACs PBS (PBS, 2% FCS) for cell counting with a hemocytometer. For cell population analysis, 1-2 million cells from cell suspensions isolated were stained 1:1000 for Aqua Live/Dead (Life Technologies), and 1:100 or 1:200 for anti-CD4 (eBioscience, CA), anti-CD8α (eBioscience), anti-CD19 (eBioscience), anti-TCRβ (eBioscience), or anti-CD45 (Biolegend, CA). Samples were analyzed with a FACSCanto (BD Biosciences) and FlowJo v10.1 (FLOWJO, OR, Ashland).

### Insulin autoantibody analyses

Frozen serum samples were analyzed for the presence of insulin autoantibodies utilizing two approaches. Cohort 1 comprised of serum from 18 GF IL10-KO mice. Cohort 1 was analyzed via radioimmunoassay previously described with an IAA cutoff based on NOD versus C57Bl/6 positive index above 10.2 (Yu *et al*., 2003). Cohort 2 comprised of serum from 11 GF IL10-KO mice and 4 SPF IL10-KO mice. Cohort 2 was analyzed via an electrochemiluminescence assay (Miao *et al*., 2015). Mouse phenotypic information and results are found in Tables 1 and 2.

**Table 1.**
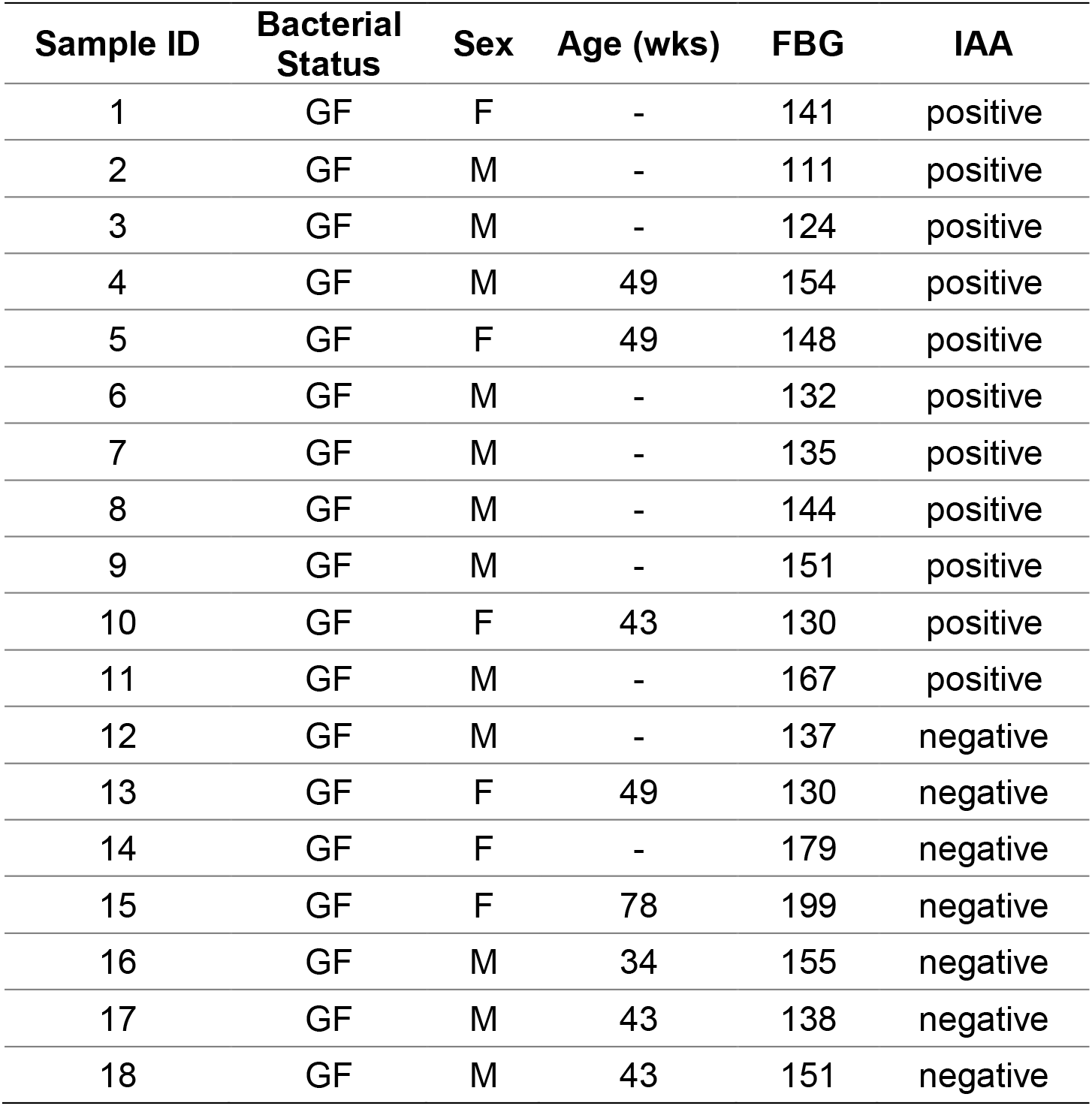
Insulin Autoantibody (IAA) testing in GF IL-10 KO Cohort 1.

**Table 2.** Insulin Autoantibody (IAA) testing in IL-10 KO Cohort 2.

| Sample ID | Bacterial Status | Sex | Age (wks) | BG* | Pancreatic Infiltration | IAA |
| --- | --- | --- | --- | --- | --- | --- |
| 1 | GF | F | 28 | 122 | positive | negative |
| 2 | GF | F | 29 | 143 | positive | negative |
| 3 | GF | F | 29 | 132 | positive | negative |
| 4 | GF | F | 31 | 199 | positive | negative |
| 5 | GF | F | 31 | 173 | positive | negative |
| 6 | GF | M | 29 | 197 | positive | negative |
| 7 | GF | M | 31 | 152 | positive | negative |
| 8 | GF | M | 34 | 159 | positive | negative |
| 9 | GF | M | 28 | 137 | negative | negative |
| 10 | GF | M | 28 | 178 | negative | negative |
| 11 | GF | M | 28 | 157 | negative | negative |
| 12 | SPF | F | 8 | - | negative | negative |
| 13 | SPF | M | 8 | - | negative | negative |
| 14 | SPF | M | 32 | *195* | negative | negative |
| 15 | SPF | M | 20 | *165* | negative | negative |

\*Blood Glucose (BG) readings are A.M. non-fasted. Italicized BG numbers indicate 4-6 hour fasted reading.

### Fecal microbiota transplantation

GF IL-10 KO mice at 4-5 weeks (“FMT-juvenile” group) and 17 weeks of age (“FMT-adult” group) were transferred from the GRAF unit into a *H. hepaticus-free* vivarium. Donor *H. hepaticus-free* mice were kindly provided by Dr. Cathryn Nagler at the University of Chicago. Fresh fecal pellets from 13-week-old WT female and male donors were diluted respectively in sterile PBS (2 pellets per 1 mL PBS). 150μL of female or male fecal slurries were administered by oral gavage into female and male recipients respectively. FMT recipient mice were monitored until 26-27 weeks of age (duration of FMT observation period 22 weeks in FMT-juvenile versus 9 weeks in FMT-adult recipients).

### Statistical analyses

Statistical analyses were performed using GraphPad Prism v7.0b via nonparametric ANOVA or unpaired t tests. Data are represented as mean ± SEM.

## Results

### GF IL-10 KO mice exhibit pancreatic infiltration characteristic of T1D histopathology

Our initial observation of polyuria in GF IL-10 KO mice prompted us to investigate whether these mice develop T1D lesions via histological assessment of pancreata. H&E stained pancreata slides revealed GF IL-10 KO mice exhibit islet and ductal lymphocytic infiltrates resembling the histopathology of T1D islet infiltrates (representative slide, Figure 1A). Immunofluorescence staining was performed to identify these immune infiltrates. Pancreatic infiltrates were positive for adaptive and innate immune markers. Infiltrates located adjacent to insulin positive β-cells were positive for CD3+ T lymphocytes, B220+ B lymphocytes, as well as CD11c+ and CD68+ macrophage and dendritic cell markers (Figure 1B-C). Interestingly, we even visualized colocalization of insulin with antigen presenting cell infiltrates (Figure 1C). We also found preliminary evidence of cytokine pathway activation with pSTAT1+ cells at the periphery of islets near infiltrates (Figure 1D).

**Figure 1.**
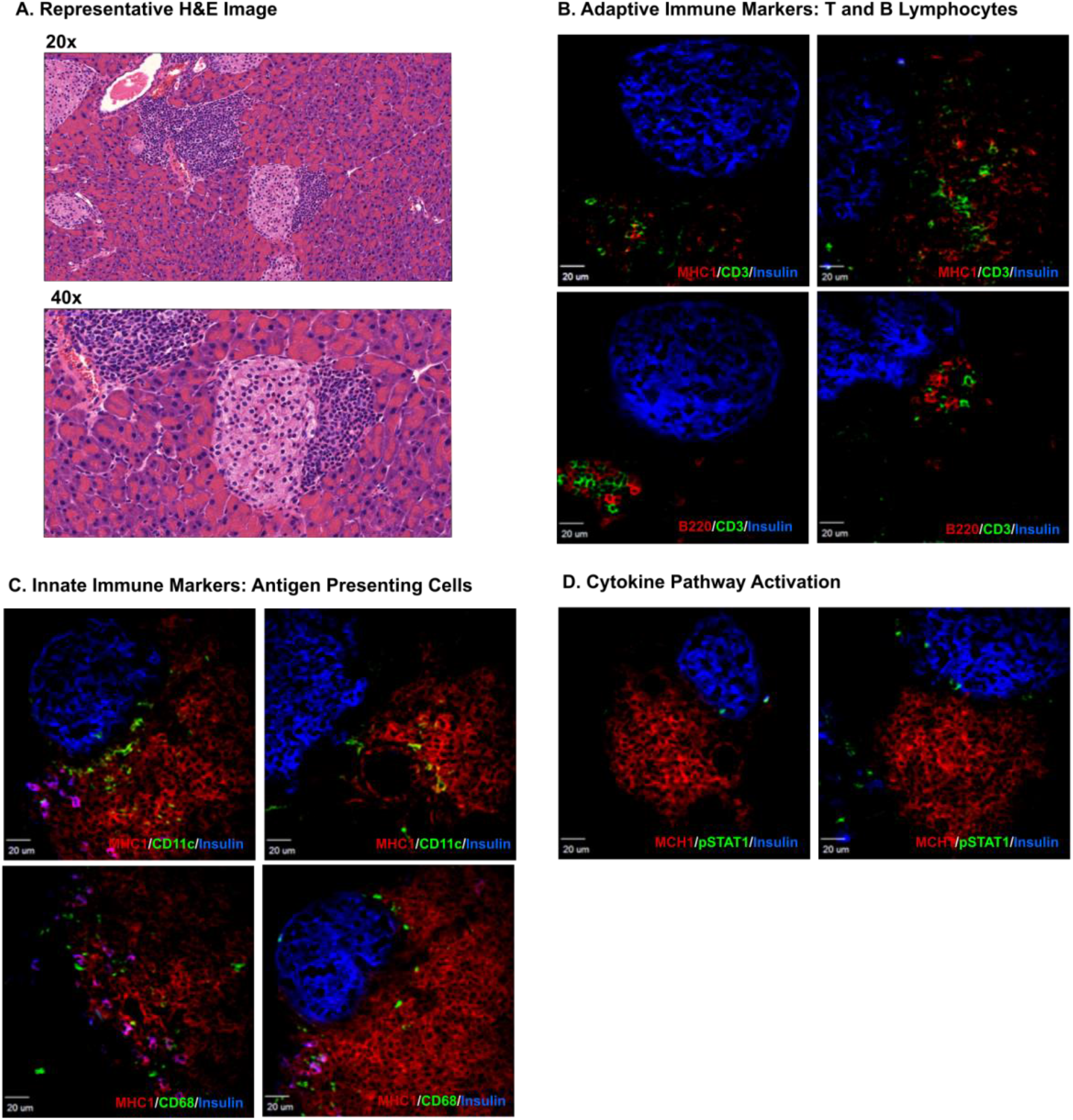
GF IL-10 KO mice develop pancreatic islet and ductal immune cell infiltrates. **A.** Representative H&E stained infiltrated pancreatic sections at 20x and 40x magnification from a GF IL-10 KO female 33 weeks of age (FBG=205mg/dL). IF staining performed on sections from two GF IL-10 KO males ~20 weeks of age reveal infiltrates are MHC class I positive (B-D). **B.** Infiltrates are positive for B220 B cell and CD3 T cell markers. **C.** Several insulin-positive cells co-stain with cells positive for CD11C or CD68, dendritic and pan-macrophage markers. **D.** Evidence of pSTAT1 suggests activation of interferon or stress-induced responses.

### Adoptive transfer of immune cells from GF IL-10 KO donors does not transfer pancreatic immunopathology into immunocompromised GF RAG1 KO recipients

A GF adoptive transfer model was employed to test the functional potential of immune cells from GF IL-10 KO mice to elicit pancreatic infiltration in GF RAG1 KO recipients that lack mature adaptive T and B lymphocytes. To determine the success of transferred immune cells, we analyzed the ratio of T and B lymphocytes in donors versus recipients. T and B lymphocytes were successfully detected via flow cytometry in recipients, but were in significantly lower frequencies as compared to donors (Figure 2A). Both female and male GF RAG1 KO recipients of IL-10 KO donors gained weight and appeared healthy throughout the 10-week observation period (Figure 2B). Likewise, wild-type (WT) donor RAG1 KO recipients appeared healthy and gained weight over the course of 9 weeks with the exception of WT-RAG1 KO females, which were older in age compared to WT-RAG1 KO recipient males (Figure 2C). Although 4/4 female donors and 1/4 male GF IL-10 KO donors were positive for pancreatic infiltration, GF RAG1 KO recipients did not exhibit pancreatic infiltration (Figure 2D). Furthermore, no differences were found in blood glucose levels between female and male recipients from either GF IL-10 KO or GF WT donors (Figures 2E and 2F).

**Figure 2.**
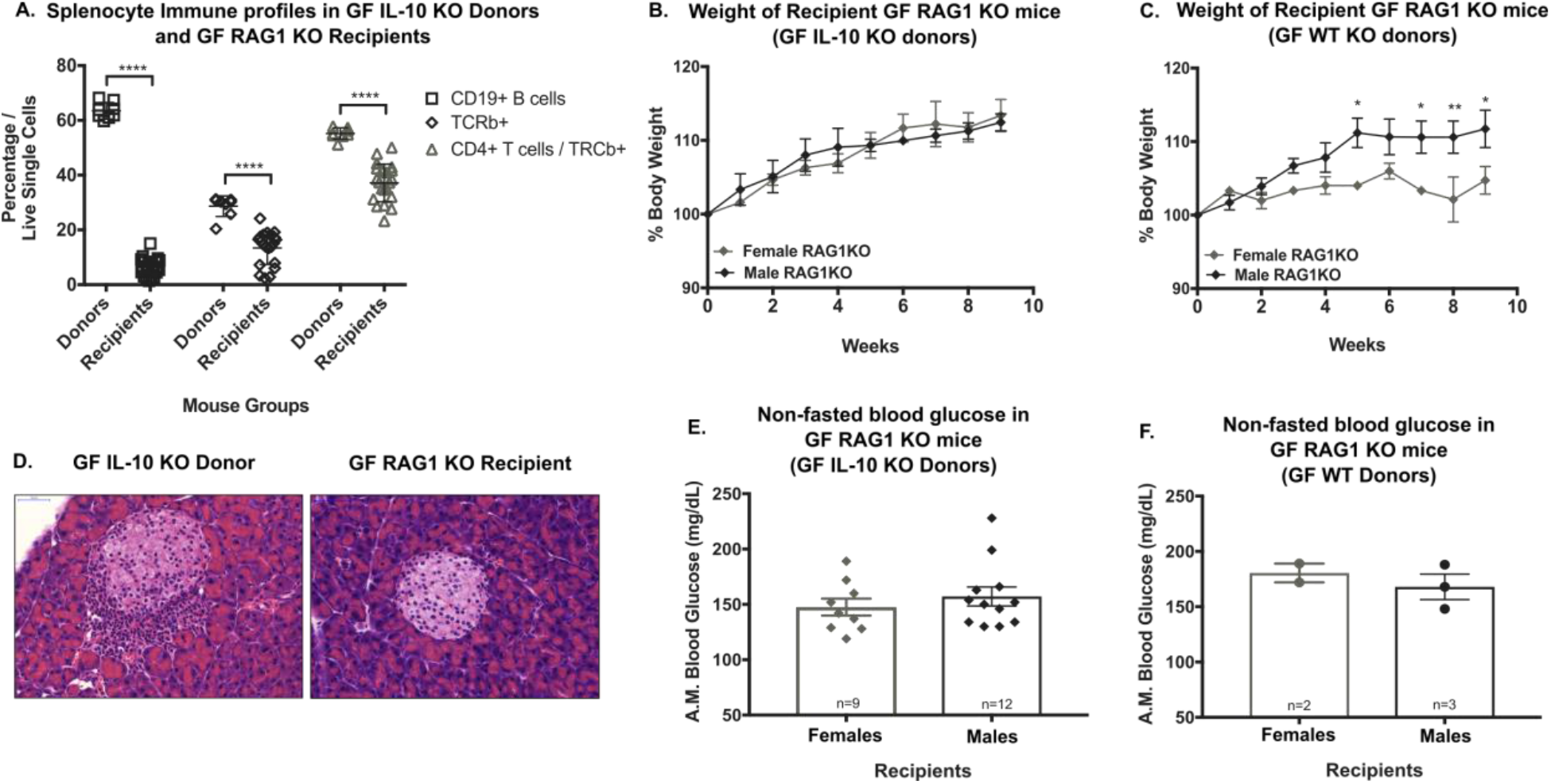
Adoptive transfer of immune cells from GF IL-10 KO donors into GF RAG1 KO recipients does not recapitulate pancreatic immunopathology. **A.** Percentage of B- and T-lymphocytes isolated from IL-10 KO donor and recipient spleens (gated live/single cell CD19+, TCRβ+ or CD4+/ TCRβ+ cells). **B.** Percent body weight of GF IL-10 KO – GF RAG1 KO female (grey diamond) and male (black diamond) recipients (n=9-12/sex). **C.** Percent body weight of GF WT – GF RAG1 KO female (grey diamond) and male (black diamond) recipients (n=2-3/sex). **D.** Representative H&E stained pancreas slide from a GF IL-10 KO female donor and a GF RAG1 KO female recipient at 40x magnification. **E.** Non-fasted A.M. blood glucose levels in female (grey diamond) versus male (black diamond) GF IL-10 KO – GF RAG1 KO recipients. **F.** Non-fasted A.M. blood glucose levels in female (grey circle) versus male (black circle) GF WT – GF RAG1 KO recipients. \*\*\*\**p*=0.0001, \*\**p*=0.006, \**p*=0.03 via Sidak’s Multiple Comparison.

### The presence of gut microbiota protects against pancreatic infiltration and GF IL-10 KO mice exhibit increased incidence of pancreatic infiltration

The presence of infiltrates resembling T1D histopathology under GF conditions led us to hypothesize (1) gut microbiota are protective against pancreatic infiltration and (2) GF IL-10 KO mice exhibit an increased incidence of pancreatic infiltration associated with T1D. We found IL-10 KO mice raised in the presence of bacteria are 100% protected against pancreatic infiltration. Conversely, 29% of GF IL-10 KO mice exhibited significant pancreatic infiltration across all ages (Figure 3A), with 50% of mice above 6-months of age positive for pancreatic infiltration (Figure 3B). This demonstrates the slow development of immunopathology in these mice. Both female and male mice displayed similar incidences, with 35% of females and 23% of males positive for pancreatic infiltration (Figure 3C). Moreover, GF WT mice displayed some signs of pancreatic infiltration in aged mice (3/10 mice >6-months of age), but this was not significant compared to the incidence found in aged GF IL-10 KO mice based on unpaired T test. This finding suggests a role for gut microbiota and IL-10 in the development of pancreatic infiltration.

**Figure 3.**
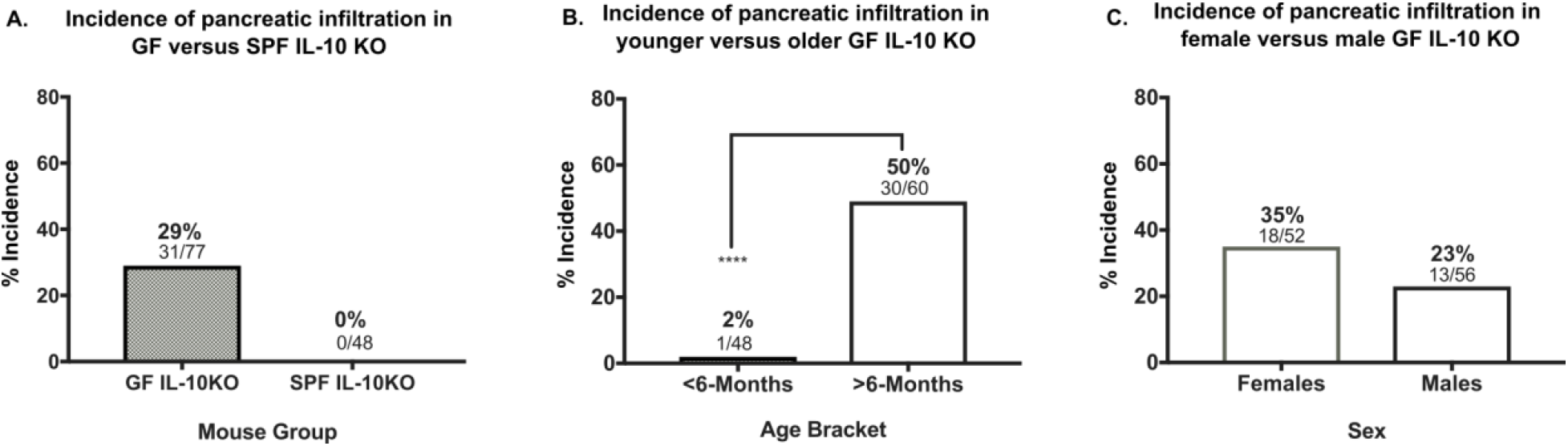
The presence of gut microbiota protects against pancreatic infiltration in IL-10 KO mice. **A.** Incidence of pancreatic infiltration in GF versus SPF IL-10 KO mice across all ages (GF n=77, SPF n=48). **B.** Incidence of pancreatic infiltration in GF IL-10 KO mice below or above 6-months of age (<6-months n=48, >6-months n=30). **C.** Incidence of pancreatic infiltration in GF IL-10 KO females versus males (female n=52, male n=56). \*\*\*\**p*>0.0001 Unpaired T test.

### A subset of GF IL-10 KO mice develop overt diabetes, but the majority of mice maintain euglycemic control

Since GF IL-10 KO mice exhibit an increased incidence for pancreatic infiltration, we tested whether this finding was associated with diabetes development, defined by blood glucose levels above 250 mg/dL. We found 4 mice, only 7% of mice above 6-months of age, developed overt diabetes. Overall, the majority of GF IL-10 KO mice did not develop overt diabetes (Figure 4A). Despite SPF IL-10 KO mice being protected from pancreatic infiltration, they displayed slightly higher levels of fasting blood glucose readings compared to age-matched GF IL-10 KO mice, but did not develop diabetes (Figure 4B). This finding was consistent with non-fasting A.M. blood glucose readings between age-matched GF and SPF IL-10 KO mice (Figure 4C).

**Figure 4.**
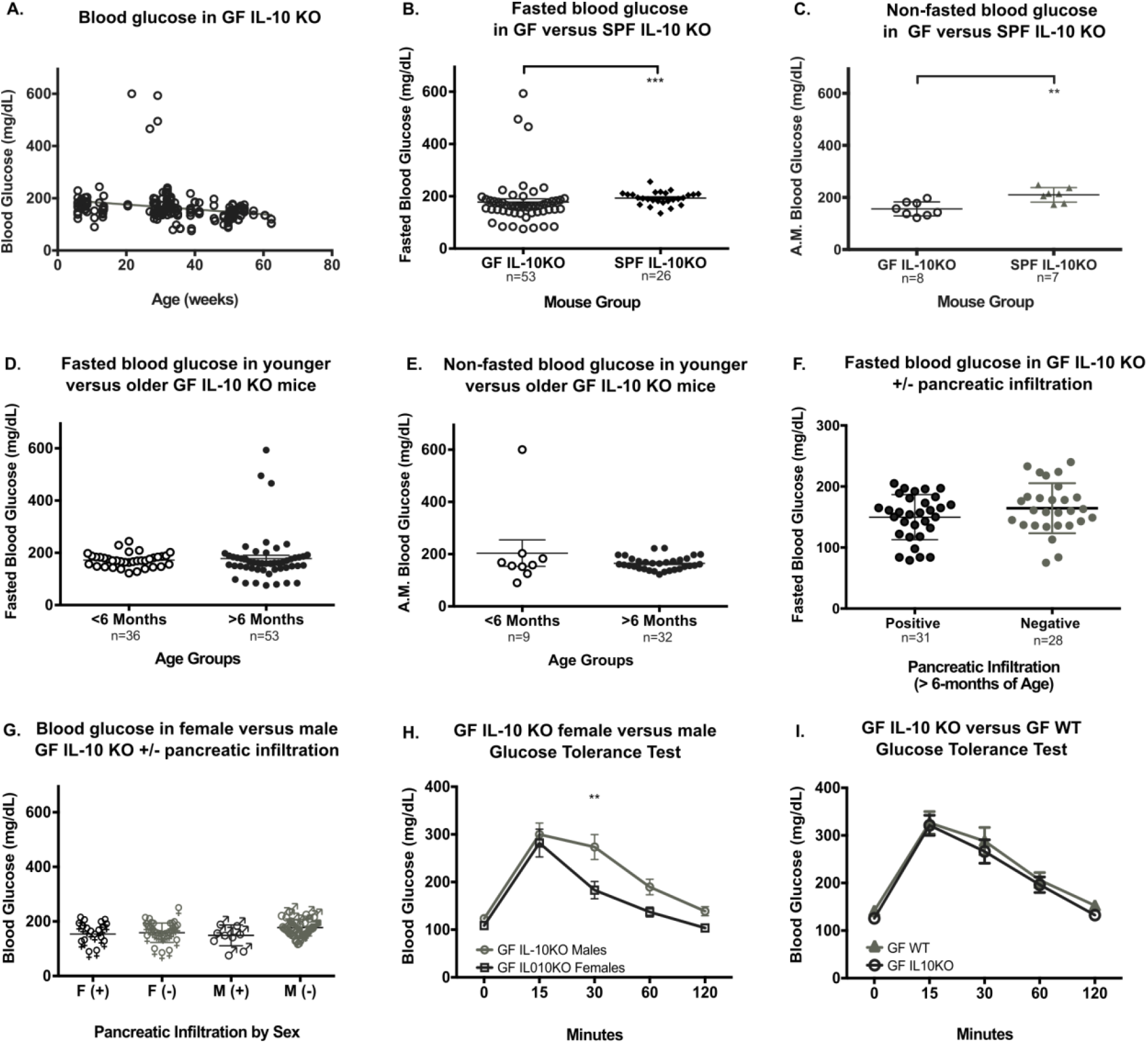
Blood glucose diabetes assessment in IL-10 KO mice. **A.** Cross-sectional blood glucose levels in GF IL-10 KO males and females (n=184). **B.** Fasted blood glucose in GF versus SPF IL-10 KO mice (clear circle, GF n=53; black diamond, SPF n=26). **C.** Non-fasted A.M. blood glucose levels in GF versus SPF IL-10 KO mice (clear circle, GF n=8; grey triangle, SPF n=7). \*\**p*=0.0024, \*\*\**p*=0.0003, via Mann-Whitney Test. **D.** Fasted blood glucose levels in GF IL-10 KO mice less than or greater than 6-months of age (clear circle, <6-months n=36; black circle, >6-months n=53). **E.** Non-fasted A.M. blood glucose levels in GF IL-10 KO mice less than or greater than 6-months of age (clear circle, <6-months n=9; black circle, >6-months n=32). **F.** Fasted blood glucose in older GF IL-10 KO mice positive or negative for pancreatic infiltration pathology (black circle, Positive n=31; clear circle, Negative n=28). **G.** Blood glucose levels in female (♀) and male (♂) GF IL-10 KO mice positive (black symbol) or negative (grey symbol) for pancreatic infiltration. **H.** Glucose tolerance test blood glucose levels in GF IL-10 KO female (black-outlined clear square) versus male (grey-outlined circle) (n=12-13/sex). **I.** Glucose tolerance test blood glucose levels in GF IL-10 KO (black-outlined circle) versus GF WT mice (grey triangle) (n=12-13/group). \*\**p*=0.002 via 2way ANOVA.

Since the incidence of pancreatic infiltration was significant in aged GF IL-10 KO mice, we compared fasting blood glucose levels in GF IL-10 KO mice above and below 6-months of age. No significant differences in fasting and non-fasting A.M. blood glucose levels were found in GF IL-10 KO mice (Figure 4D-E). We next tested the hypothesis that GF IL-10KO mice with pancreatic lymphocytic infiltrates resembling the pathology of T1D would have higher fasting blood glucose levels compared to mice without infiltration. Despite differences in the presence or absence of pancreatic immunopathology, age-matched GF IL-10 KO mice did not exhibit significant differences in fasting blood glucose levels (Figure 4F). Additionally, no gender dimorphism was observed in blood glucose levels between female and male GF IL-10 KO mice (Figure 4G). Although no significant differences in blood glucose levels were observed between sexes, glucose tolerance tests revealed differences in glucose tolerance between older GF IL-10 KO females and males. GF IL-10 KO females appear to be slightly more glucose tolerant compared to males (Figure 4H). No differences in glucose tolerance were found in GF WT females versus males (Figure 4I).

### A subset of GF IL-10 KO mice are positive for insulin autoantibodies

Since the majority of mice maintain glycemia and IF staining revealed co-localization of immune cell infiltrates with insulin, we investigated whether this model was analogous to stage 1 of T1D in humans, in which patients are euglycemic and positive for the presence of autoantibodies (Insel *et al*., 2015). We hypothesized that GF IL-10 KO mice would be positive for insulin autoantibodies (IAA), one of the first autoantibodies to appear in pre-diabetic patients (Ziegler *et al*., 1993). We therefore tested for the presence of IAA in the sera of GF IL-10 KO mice originally suspected of diabetes (Cohort 1, Table 1). These mice exhibited varying degrees of pancreatic infiltration and had elevated fasting blood glucose (FBG) levels but were not diabetic. In Cohort 1, 61% (11/18 mice) were positive for IAA. We also tested a second cohort of non-fasted and fasted mice (Cohort 2, Table 2), which included 11 GF IL-10 KO mice above 6-months of age that were positive or negative for pancreatic infiltration. However, none of the mice in Cohort 2 were positive for IAA.

### The reintroduction of gut microbiota via fecal microbiota transplantation protects IL-10 KO recipients against pancreatic infiltration and does not impact blood glucose levels

Fecal microbiota transplantation (FMT) has become a useful tool for patients clinically and in research as a means to study the role of gut microbiota in disease pathogenesis in the lab. We utilized FMT during juvenility and adulthood to test the role of gut microbiota in the protection against pancreatic infiltration in GF IL-10 KO recipients. We addressed two factors that could impact our ability to sufficiently monitor mice for diabetes development: FMT recipient age and the presence of the pathobiont, *H. hepaticus* (Kullberg *et al*., 1998). To assess pancreatic immunopathology and T1D development, *H. hepaticus*-free donor gut microbiota was introduced into GF IL-10 KO females and males at 5 weeks (FMT-juvenile) or 17 weeks of age (FMT-adult). We hypothesized FMT would be more protective against pancreatic infiltration when introduced early in life during juvenility as compared to adulthood. FMT IL-10 KO recipients in both age groups appeared to be protected from pancreatic infiltration (Figure 5A). Similar to our blood glucose assessment in SPF IL-10 KO mice, FMT IL-10 KO mice had slightly higher non-fasting A.M. blood glucose levels compared to age-matched GF IL-10 KO mice (Figure 5B). Moreover, age of bacterial introduction into juvenile versus adult recipients did not impact overall blood glucose levels in age-matched FMT IL-10 KO recipients at time of sacrifice, 27 weeks of age (Figure 5C). Likewise, glucose tolerance test performed 8-11 weeks post-FMT demonstrated no difference in glucose tolerance between FMT-juvenile versus FMT-adult IL-10 KO and WT mice (Figure 5D and 5E).

**Figure 5.**
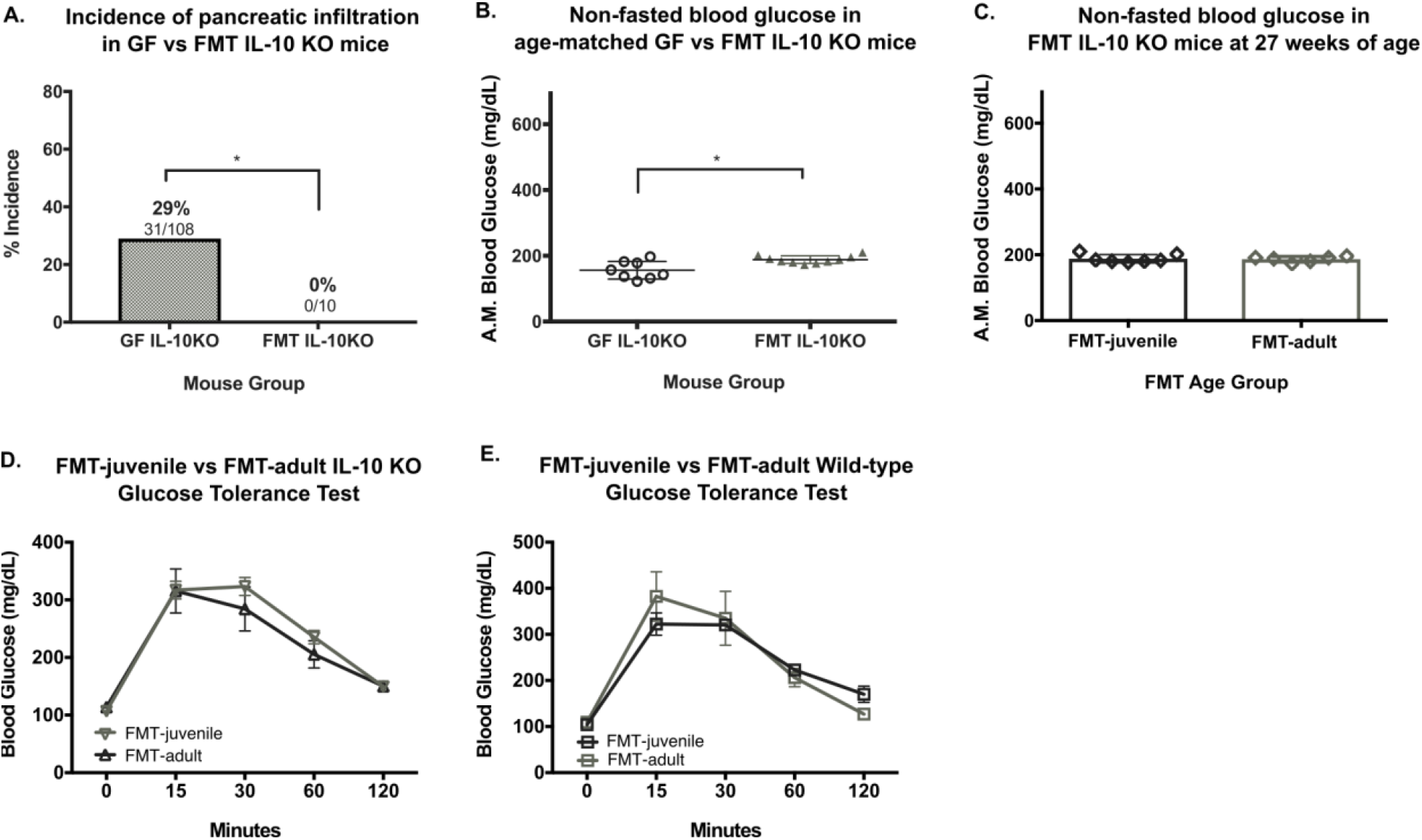
Fecal microbiota transplantation (FMT) of GF IL-10 KO recipients protects against pancreatic infiltration and does not impact blood glucose levels of FMT recipients. **A.** Incidence of pancreatic infiltration in GF versus FMT IL-10 KO mice (GF n=108, FMT n=10) \**p*=0.049 Unpaired T Test. **B.** Non-fasted A.M. blood glucose levels in age-matched GF (clear circle) versus FMT (grey triangle) IL-10 KO mice (n=8-10/group) \**p*=0.015 via Mann-Whitney Test. **C.** Non-fasted A.M. blood glucose levels in FMT-juvenile (black-outlined diamond) versus FMT-adult (grey-outlined diamond) IL-10 KO mice at 27 weeks of age (n=6-7/group). **D.** Glucose tolerance blood glucose levels in FMT-juvenile (grey-outlined upside-down-triangle) versus FMT-adult (black-outlined triangle) IL-10 KO mice (n=6-7/group). **E.** Glucose tolerance blood glucose levels in FMT-juvenile (black-outlined square) versus FMT-adult (grey-outlined square) wild-type mice (n=3-4/group).

## Discussion

The present work reveals two novel findings: first, GF IL-10 KO mice exhibit early phases of pancreatic immune infiltration resembling T1D histopathology, and second, the presence of gut microbiota in SPF IL-10 KO mice is protective against this phenomenon (Bobe *et al*., 2017). This model addresses some current limitations to the field by allowing us to study disease initiation and increase our understanding of microbe-mediated immune mechanisms underlying disease. The use of GF IL-10 KO mice to initially study the role of gut microbiome in inflammatory bowel diseases led us to the discovery of pancreatic infiltration and diabetes in a subset of aged GF IL-10 KO mice. We hypothesized that the loss of microbial signals and IL-10 play a protective role in the development of pancreatic immunopathology. We focused on immune-histopathology, blood glucose assessment, and an adoptive transfer model to test whether GF IL-10 KO exhibit an increased incidence of pancreatic infiltration associated with T1D. We then determined whether gut microbiota is protective and utilized a FMT model to assess the pancreas of FMT IL-10 KO mice versus SPF IL-10 KO counterparts.

Pancreatic data from T1D donors and murine models support the role of B and T lymphocytes and antigen-presenting cells in disease pathology (In’t Veld, 2011; Magnuson *et al*., 2014; Walker and von Herrath, 2016). Our immunofluorescence staining of pancreatic infiltrates in GF IL-10 KO mice confirms the presence of T and B lymphoid cells and myeloid cells, such as macrophages and dendritic cells. We also observed positive pSTAT1 staining at the periphery of islets near infiltrates, which has been implicated in T1D pathogenesis and β-cell death (Sunshin Kim et al., 2007). Hence, our immunostaining results are compatible with previous human and mouse staining of islet infiltrates (Willcox *et al*., 2009; Coppieters *et al*., 2012; Thompson *et al*., 2014; Carrero, Ferris and Unanue, 2016; Lundberg *et al*., 2017). We characterized the immunogenicity of GF IL-10 KO mice via adoptive transfer of splenic immune cells into immunodeficient GF RAG1 KO recipients lacking mature T and B lymphocytes (Mombaerts *et al*., 1992). The failure to propagate the disease phenotype in recipient mice may be contributed to donors exhibiting only mild pancreatic infiltration and no diabetes. The frequency of transferred T and B cells in recipients was also significantly lower in spleens of recipients (Figure 2), thus, transferred donor cells may not have lived long enough or migrated to the pancreatic lymph nodes and pancreas to propagate disease pathology.

Interestingly, the majority of GF IL-10 KO mice maintained euglycemia and only a small subset of mice became overtly diabetic. The delayed onset of pancreatic immunopathology and diabetes may in part be due to the C57Bl/6 genetic background and MHC b haplotype, which is thought to be more protective against diabetes compared to the g7 haplotype found in the diabetes-prone non-obese diabetic (NOD) mouse model (Leiter, Coleman and Hummel, 1981). While genetic analyses of GF IL-10 KO mice were beyond the scope of this study, this model offers an intriguing opportunity to refine genetic and environmental factors that dictate the natural history of disease development. A modified disease classification system has been proposed to capture the sequential nature of disease starting from presymptomatic to symptomatic stages (Insel *et al*., 2015). Our model reflects the first of three progressive stages in T1D development in which patients maintain euglycemia but are positive for autoantibodies. The presence of autoantibodies specific to islets before disease diagnosis has been a hallmark of human T1D, with insulin as one of the major targets of humoral autoimmunity in humans and the NOD model (Bonifacio *et al*., 2001; Ziegler *et al*., 2011). Greater knowledge on autoantibodies and autoreactive cells in this model would allow for more sophisticated techniques to uncover immune mechanisms involved in this model.

Significantly higher pancreatic infiltration in both female and male GF IL-10 KO mice at an older age demonstrates a slow and progressive nature of disease, which may require additional triggers to accelerate to overt diabetes. This finding also suggests age plays a role in disease development in the absence of specific microbial signals and potentially IL-10 mediated immunoregulatory mechanisms (Thompson *et al*., 2014). Current knowledge and outstanding questions regarding the immune mechanisms involved in the development of autoimmune T1D have been previously reviewed (Wå and Cooke, 2013; Christoffersson and von Herrath, 2016; Pugliese *et al*., 2016; Tai, Wong and Wen, 2016; Walker and von Herrath, 2016). One possible explanation for the pancreatic phenotype in our mice is immune dysregulation due to the lack of IL-10 and microbial-mediated signals. IL-10 signaling is important for Treg cell function in restraining T cell-mediated inflammation (Chaudhry *et al*., 2011; Keubler *et al*., 2015). When Foxp3-expressing Treg cells are ablated, conventionally-raised mice develop widespread immune-mediated lesions (Kim, Rasmussen and Rudensky, 2007; Chinen *et al*., 2010). Treg cells also regulate diabetogenesis in NOD mice, with a temporal decline in functional potency (Tritt *et al*., 2008). Furthermore, GF conditions may potentiate immunogenicity because mice raised GF contain fewer Treg cells compared to conventionally-raised mice (Östman *et al*., 2006).

Although the absence of microbial signals in GF IL-10 KO prevents the development of spontaneous colitis (Sellon *et al*., 1998), the loss of potentially protective microbial-mediated signals may contribute to the immunopathology we observe in the pancreas. A large body of literature supports microbial dysbiosis as a major contributor to disease pathogenesis (Vaarala, 2012; Gulden, Wong and Wen, 2015; Knip and Siljander, 2016; Paun, Yau and Danska, 2016; Hu, Wong and Wen, 2017). Work from our lab also demonstrates disturbances to gut microbial communities early in life impacts the immune system of IL-10 KO mice and increases IBD incidence (Miyoshi *et al*., 2017). Our findings here support the notion that microbial manipulation may lower risk of disease in genetically susceptible individuals. We hypothesized that exposure to microbial signals at an early age would be more protective against pancreatic infiltration compared to exposure in adulthood. Interestingly, both juvenile (5 weeks of age) and adult (17 weeks of age) FMT IL-10 KO recipients were protected from pancreatic infiltration. However, a subset of mice developed colitis, with FMT-adult mice developing more severe colitis more rapidly than FMT-juvenile mice (data not shown). These preliminary results support a role for microbial signals in immune regulatory mechanisms presumably via connections between the gut and the pancreas (Turley *et al*., 2005; Paun, Yau and Danska, 2016). FMT may stimulate microbe-mediated immune signaling pathways that direct inflammation in the gut rather than the pancreas of IL-10 KO mice. Further assessment of gut integrity, microbial assemblage, and microbe-mediated immune signaling pathways would contribute to the growing understanding of potential microbial-based therapies for complex immune disorders such as IBD and T1D (Wu and Wu, 2012; Gulden, Wong and Wen, 2015).

Since diabetes is halted at an early stage of T1D pathogenesis in our GF IL-10 KO mouse model, further investigation into immunological and metabolic pathways at play would yield more mechanistic insights and contribute to potential prophylactic treatment strategies. Moreover, studies focused on metabolic mechanisms underlying T1D pathogenesis suggest metabolomic disturbances precede the onset of T1D (Mejía-León and Barca, 2015; Overgaard *et al*., 2016). Thus, future investigations could focus on specific environmental triggers capable of fueling metabolic and immune dysfunction, particularly through cellular mechanisms that originate in the gut and impact host immune and metabolic responses. For example, postprandial nutrients and microbial signals impact enteroendocrine cells, such as L cells, which secrete hormones like glucagon-like peptide-1 (GLP-1) (Greiner and Bäckhed, 2016). GLP-1 and GLP-1 receptors regulate several physiological functions in both the pancreas and the gut. In addition to its role as an incretin, GLP-1 signaling also modulates innate immune responses in the gut via enteroendocrine-immune interactions (Yusta *et al*., 2015). Interestingly, GF mice exhibit elevated GLP-1 levels, as well as altered metabolism (Wichmann *et al*., 2013; Greiner and Bäckhed, 2016). Thus, elevated GLP-1 and other enteroendocrine hormones may participate in driving immune responsiveness and immunopathology in the pancreas of GF IL-10 KO mice. Future investigation of both systemic and more localized physiologically changes in the gut and the pancreas may reveal mechanisms perpetuating autoimmune responses. Characterization of metabolic profiles in our model would shed light on the cross talk between the enteroendocrine system, immunity, and metabolism.

In summary, our initial characterization of pancreatic infiltration in IL-10 KO mice provides the foundation to understanding the role of gut microbiota in early stages of T1D disease immunopathology. The IL-10 KO mouse model allows for the investigation of gut-mediated signaling in disease pathogenesis of the intestine and the pancreas in the presence and absence of the microbiome. While further characterization is required to dissect specific immune compartments and the role of microbial-mediated signaling pathways, our findings support the notion that targeting the microbiome and/or microbial-mediated immune signaling pathways could treat, prevent, or even reverse disease (Kondrashova and Hyöty, 2014; Burrows *et al*., 2015; Itoh and Ridgway, 2017). Our work and the amalgamation of knowledge from future studies will perhaps conceive the development of microbe-mediated prophylactic strategies for patients (Gulden, Wong and Wen, 2015; Garyu, Meffre and Cotsapas, 2016). Thus, manipulation of the microbiome may one day lower diabetes risk in genetically prone individuals.

## Acknowledgements

We thank the University of Chicago GRAF staff for their diligent work with our GF mice (Michael Yabes, Megan Bales, Melanie Spedale, and Christina Olivares), the HTRC Core facility for histological advice and processing (Terri Li, Christy Schmehl, Xin Jiang, and Ming Cheng), Dr. Alexander (Sasha) Chervonsky, University of Chicago, for his immunological and T1D animal model counsel, and Dr. Lou Philipson for his input on β-cells and diabetes. We also thank Jason B. Williams, University of Chicago Gajewski lab, for his guidance with flow cytometry, the laboratory of Dr. Bana Jabri for sharing flow cytometry antibodies, and Dr. Amanda Pasgai, University of Florida Atkinson Lab, for her feedback and help in reviewing this manuscript.

